# An Analysis of Decision Under Risk in Rats

**DOI:** 10.1101/446575

**Authors:** Christine M. Constantinople, Alex T. Piet, Carlos D. Brody

## Abstract

Prospect Theory is the predominant behavioral economic theory describing decision-making under risk. It accounts for near universal aspects of human choice behavior whose prevalence may reflect fundamental neural mechanisms. We now apply Prospect Theory’s framework to rodents, using a task in which rats chose between guaranteed and probabilistic rewards. Like humans, rats distorted probabilities and showed diminishing marginal sensitivity, in which they were less sensitive to differences in larger rewards. They exhibited reference dependence, in which the valence of outcomes (gain or loss) was determined by an internal reference point reflecting reward history. The similarities between rats and humans suggest conserved neural substrates, and enable application of powerful molecular/circuit tools to study mechanisms of psychological phenomena from behavioral economics.

---

In 1979, Daniel Kahneman and Amos Tversky published a ground-breaking paper titled “Prospect Theory: An Analysis of Decision Under Risk,” which presented a behavioral economic theory that could account for the ways in which humans deviate from normative (i.e., optimal) behavior (1, 2). For example, people exhibit probability distortion (they overweight low probabilities), loss aversion (losses loom larger than gains), and reference dependence (outcomes are evaluated as gains or losses relative to an internal reference point; fig. S1). These phenomena have been observed in laboratory and economic settings (3). Policies in the public and private sectors have been implemented, such as automatic enrollment and increases in contributions to retirement savings, to counteract these phenomena and promote smarter fiscal behavior (4, 5).

Most behavioral economic studies examine decisions between clearly described probabilistic outcomes (i.e., “decisions from description”). Studies of risky choice in rodents, however, typically examine decisions between prospects that are learned over time (i.e., “decisions from experience”), which can produce qualitatively different behavior (6). We designed a task in which the reward and probability of prospects are communicated by sensory evidence trial-by-trial, eliciting decisions from description rather than experience. This allowed us to apply core behavioral economic approaches in rodents for the first time.

Rats initiated a trial by nose-poking in the center port of a three-port wall. Light flashes were presented from left and right side ports, and the number of flashes conveyed the probability of receiving water reward at each port. Simultaneously, auditory clicks were presented from speakers on the left and right sides, and the click rate conveyed the volume of water reward baited at each port (Fig. 1A, B). One port offered a guaranteed or “safe” reward, and the other offered a risky reward with an explicitly cued probability. The safe and risky ports (left/right) varied randomly on each trial. One of four water volumes could be the guaranteed or risky reward (6, 12, 24, 48μL), and the risky reward probabilities ranged from 0 to 1, in increments of 0.1 (Fig. 1A, B).

High-throughput behavioral training generated many trained rats (n=36) and many tens of thousands of choices per rat, enabling detailed quantification of behavior. Rats demonstrated they learned the meaning of the cues by frequently “opting-out” of trials offering smaller rewards, leaving the center poke despite incurring a time-out penalty and white-noise sound (fig. S2). This indicated that they associated the click rates with water volumes, instead of using a purely perceptual heuristic. Rats favored the prospect with the higher expected value (volume x probability; Fig. 1C, fig. S2).

**Fig. 1.**
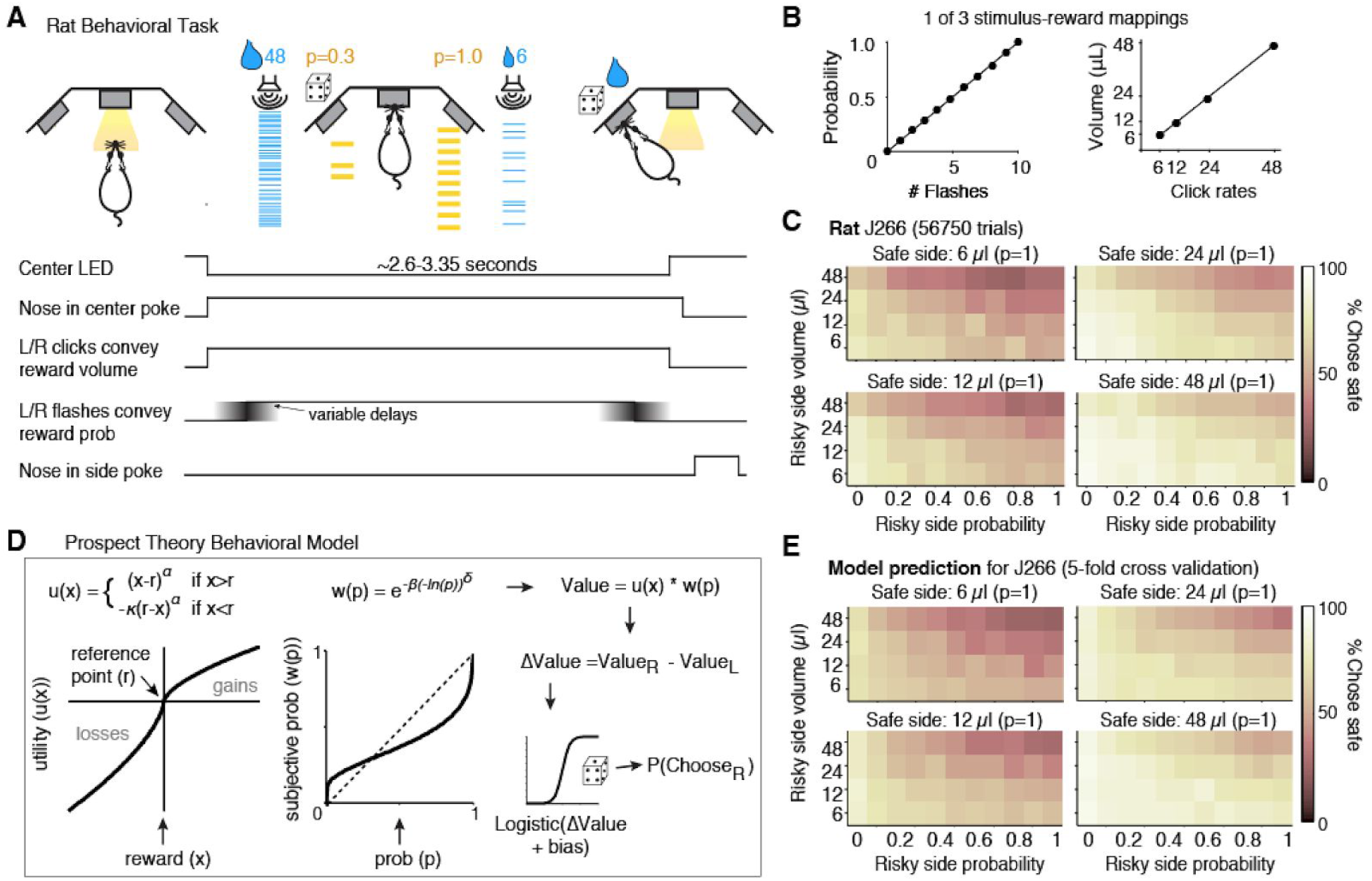
Rats choose between guaranteed and probabilistic rewards. **(A)** Behavioral task and timing of task events: flashes cue reward probability (p) and click rates convey water volume (x) on each side. Safe and risky sides are not fixed. **(B)** Relationship between cues and reward probability/volume in one task version. Alternative versions produced similar results (fig. S4). There were four possible volumes (6, 12, 24, or 48μL), and the risky side offered reward probabilities between 0 and 1 in increments of 0.1. **(C)** One rat’s performance for each of the safe side volumes. Axes are probability and volume of risky options. **(D)** A behavioral model inferred the utility and probability weighting functions that best explained rats’ choices. We modeled the probability that the rat chose the right side by a logistic function whose argument was the difference between the subjective value of each option (*V_R_-V_L_*) plus a trial history-dependent term. Subjective utility was parameterized as:

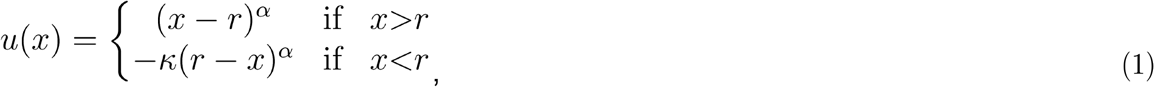

where *α* is a free parameter, and *x* is reward volume. *r* is the reference point, which determines whether rewards are perceived as gains or losses. We first consider the case where *r* = 0, so *u*(*x*) = *x^α^*. The subjective probability of each option is computed by:

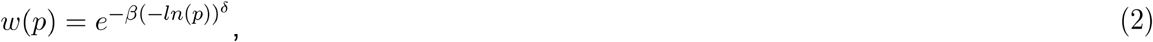

where *β* and *δ* are free parameters and *p* is the objective probability offered. Combining utility and probability yields the subjective value for each option:

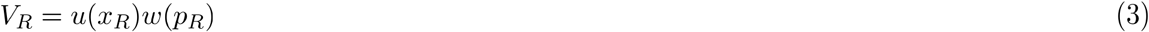

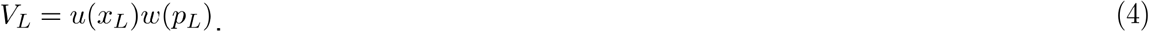 These were transformed into choice probabilities via a logistic function:

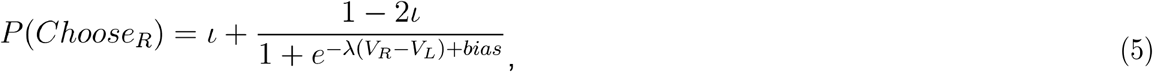

where 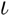 captures stimulus independent variability (lapse rate) and λ determines the sensitivity of choices to the difference in subjective value (*V_R_ – V_L_*). The bias term was comprised of three possible parameters, depending on trial history:

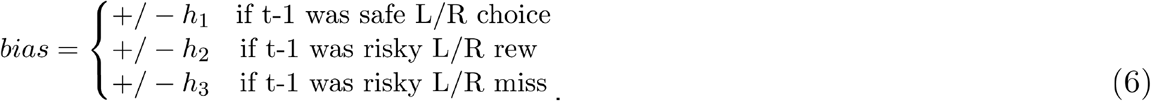 **(E)** Model prediction for held-out data from one rat, averaged over 5 test sets.

Two key economic concepts are *utility* (rewards are evaluated by the subjective satisfaction or “utility” they provide) and *probability distortion* (people often overweight low probabilities; Fig. 1D; fig. S1). We used a standard choice model (7, 8) to estimate each rat’s utility and probability weighting functions according to Prospect Theory (Fig. 1D; see legend). The model predicted rats’ choices on held-out data (Fig. 1E). It outperformed alternative models, including one imposing linear probability weighting (according to Expected Utility Theory (9)), and one that fit linear weights for probabilities and volumes (fig. S3).

Concave utility (the utility function exponent *α*<1) produces diminishing marginal sensitivity, in which subjects are less sensitive to differences in larger rewards. Kahneman and Tversky (2) observed a median *α* of 0.88 in humans; the median *α* for our rats was 0.54, consistent with concave utility, as in humans (Fig. 2A). To test for diminishing marginal sensitivity, we compared rats’ performance when choosing between guaranteed outcomes of 0 or 24μL, and 24 or 48μL (Fig. 2B, C). For rats with more concave utility functions, 24 and 48μL should be less discriminable than 0 and 24μL, leading to worse performance on these trials (Fig. 2C). Indeed, the concavity of the utility function was significantly correlated with reduced discriminability on trials offering 24 and 48μL (Fig. 2D, E; p = 1.03e-7, Pearson’s correlation). Rats, like humans, exhibit diminishing marginal sensitivity.

Model fits for rats’ probability weighting functions revealed overweighting of probabilities (Fig. 2F). A logistic regression model that parameterized each probability to predict choice yielded regression weights that mirrored the probability weighting functions (Fig. 2G, H). Control experiments indicated that nonlinear utility and probability weighting were not due to perceptual errors in estimating the numbers of flashes and clicks (10) (fig. S4).

**Fig. 2.**
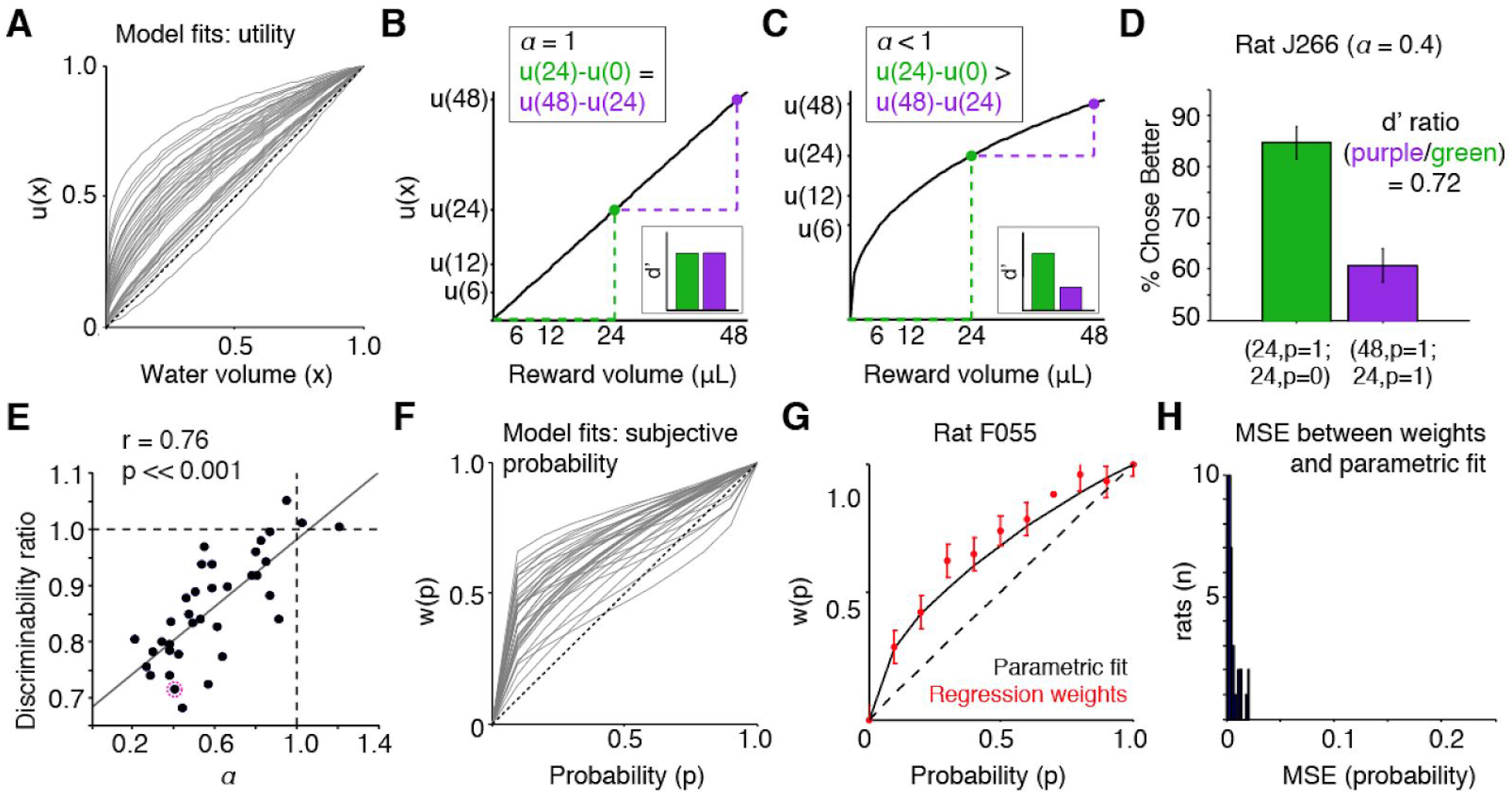
Non-parametric analyses confirm nonlinear utility and probability weighting. **(A)** Model fits of subjective utility functions for each rat, normalized by the maximum volume (48μL). **(B)** Schematic linear utility function: the perceptual distance (or discriminability, d’) between 0μL and 24μL is the same as 24μL and 48μL. **(C)** Schematic concave utility function: 24μL and 48μL are less discriminable than 0μL and 24μL. **(D)** One rat’s performance on trials with guaranteed outcomes of 0μL vs. 24μL (green), or 24μL vs. 48μL (purple). Performance ratio on these trials (“d’ ratio”) less than 1 indicates diminishing sensitivity. **(E)** The concavity of the utility function (*α*) is significantly correlated with reduced discriminability of larger rewards. Pink circle is rat from D. **(F)** Model fits of probability weighting functions. **(G)** Weights from logistic regression parameterizing each probability match probability weighting function for one rat. Error bars are s.e.m. for each regression coefficient. **(H)** Mean squared error between regression weights and parametric fits for each rat (mean mse=0.006, in units of probability).

To evaluate rats’ risk attitudes, we used a non-parametric assay and measured the *certainty equivalents* for all gambles of 48μL (2, 11, 12). The certainty equivalent is the guaranteed reward the rat indicates is of equal value to the gamble (Fig. 3A, B). If it is less than the gamble’s expected value, the subject is risk averse: he effectively “undervalues” the gamble, and is willing to accept a smaller guaranteed reward to avoid risk (Fig. 3C). Conversely, if the certainty equivalent is greater than the gamble’s expected value, the subject is risk seeking (Fig. 3D), and risk neutral if they are equal. Measured certainty equivalents closely matched those predicted from the model, using an analytic expression incorporating utility and probability weighting functions (*CE* = *w*(*p*)^1/*α*^; Methods. Pearson’s correlation 0.96, p=1.58e-11; Fig. 3C, D). This non-parametric assay further validated the model fits and revealed heterogeneous risk preferences across rats (Fig. 3E; fig. S5).

**Fig. 3.**
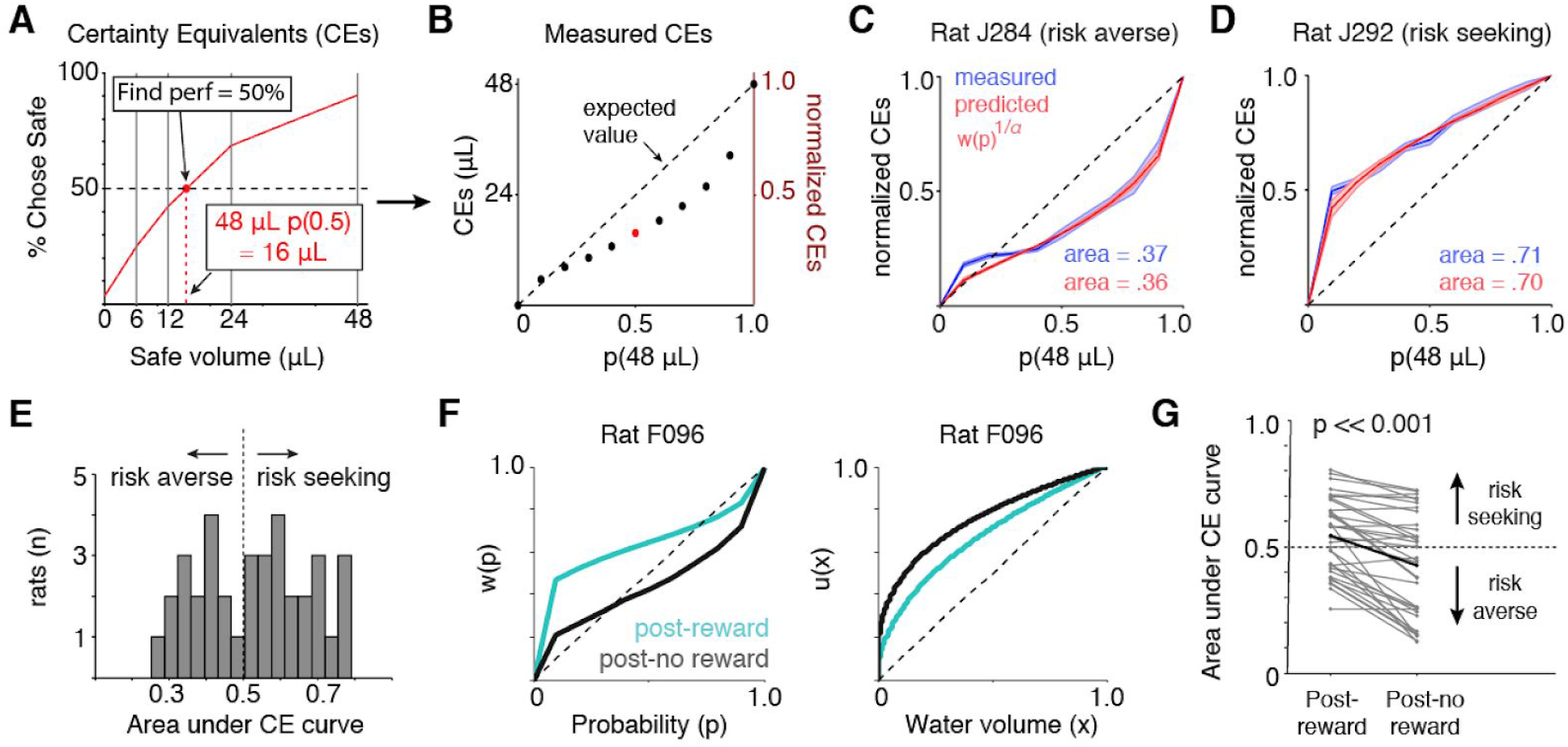
Rats exhibit diverse risk attitudes that are modified by reward history. **(A, B)** To obtain “certainty equivalents,” we measured psychometric functions for each probability of receiving 48μL, and estimated the certain volume at which performance = 50%. **(C)** Measured (blue) and model-predicted (red) certainty equivalents from one rat indicates systematic undervaluing of the gamble, or risk aversion. Error bars for model-prediction are 95% confidence intervals of parameters from 5-fold cross validation. Data are mean +/- s.e.m. for left-out test sets. **(D)** Same as C for a risk-seeking rat. **(E)** Distribution of CE areas computed using analytic expression from model fits. **(F)** Probability weighting function (left) and utility function (right) for one rat from model fit to trials following reward (turquoise) or no reward (black). **(G)** CE areas predicted from model fits for all rats following rewarded and unrewarded trials.

Risk attitudes are assumed to be stable traits, but may be flexible; for example, stock returns affect investors’ subsequent risk aversion (13, 14). Fitting the model separately to trials following rewarded and unrewarded choices revealed systematic shifts in the utility and probability weighting functions: utility functions became less concave and probability weighting functions became more elevated to reflect increased likelihood of risky choice following rewards (Fig. 3F). This was consistent across rats, as observed in the model-predicted certainty equivalents (Fig. 3G).

Another key feature of human choice behavior is “reference dependence:” people evaluate rewards as gains or losses relative to an internal reference point. It is unclear what determines the reference point (3); proposals include status quo wealth (1), reward expectation (15, 16), heuristics based on the prospects themselves (17, 18), or recent reward experience (19, 20).

**Fig. 4.**
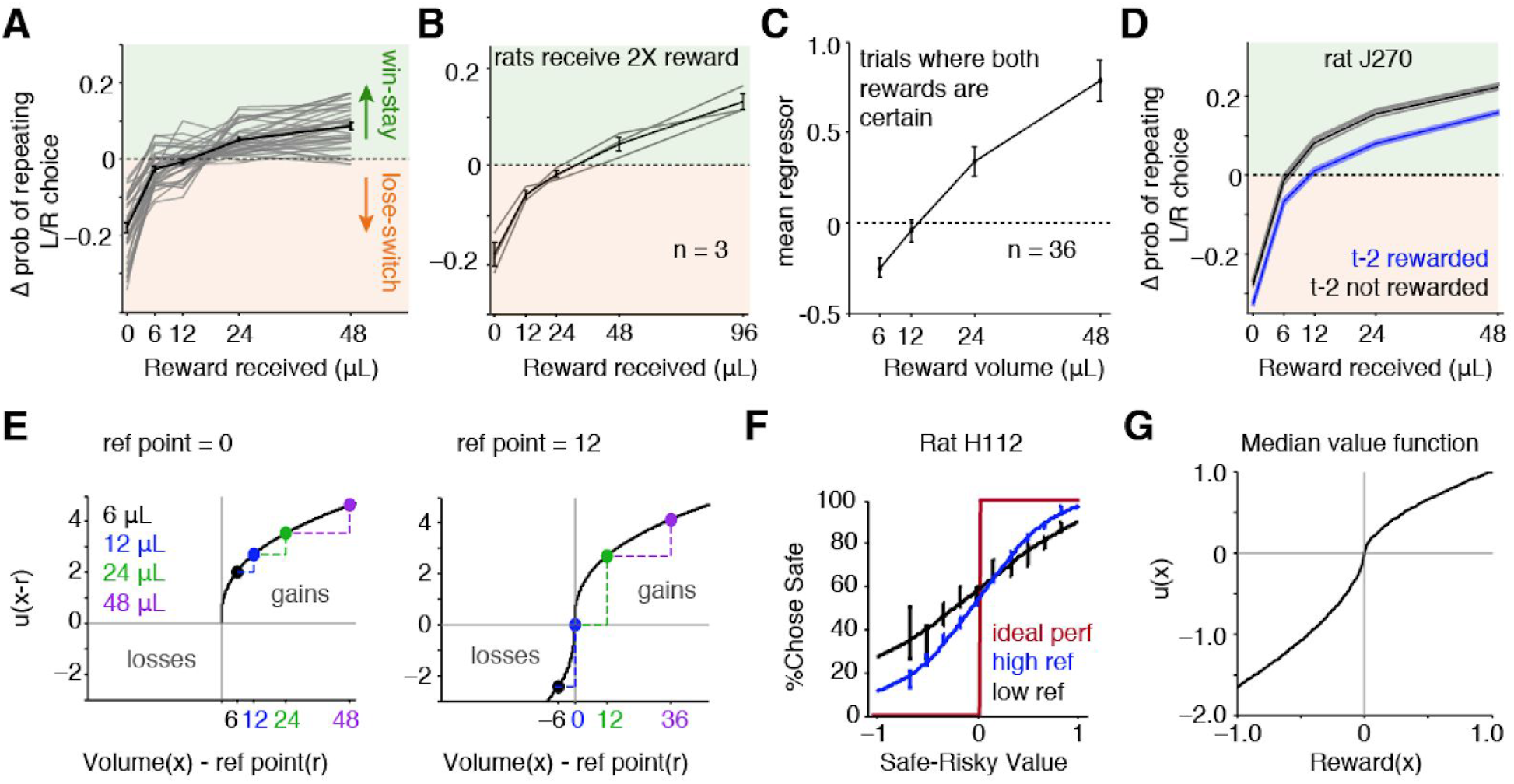
Rats exhibit reference dependence. **(A)** ΔProbability of repeating left/right choices (relative to mean probability of repeating), following each reward. Points above the dashed line indicate an increased probability of repeating (“stay”); those below indicate a decreased probability (“switch”). Black curve is average +/− s.e.m. across rats. **(B)** 3 rats were trained with doubled water volumes. They exhibited lose-switch biases following 12 and 24μL. **(C)** A logistic regression model predicted rats’ choices based on offered reward volumes, on trials when both sides offered guaranteed rewards. On average, weights for 6μL were negative. **(D)** Win-stay/lose-switch biases for one rat separated by reward history two trials back. **(E)** Schematic illustrating that with concave utility, rewards should be more (less) discriminable when the reference point is high (low). **(F)** Psychometric performance from one rat when the inferred reference point was low (black) or high (blue). Red curve is ideal performance. **(G)** Value function with the median parameters across rats indicates loss aversion (median *α*=0.6, *k*=1.7).

Rats exhibited reference dependence by treating small rewards (6/12μL) as losses. They exhibited win-stay/lose-switch biases: following unrewarded trials, rats were more likely to switch ports (Fig. 4A). Surprisingly, most rats exhibited a “switch” bias after receiving 6 or 12μL. Reward can only be considered a loss relative to a reference point. The “win/lose” threshold (i.e., reference point) was experience-dependent: 3 rats trained with doubled reward volumes (12-96μL) exhibited lose-switch biases after receiving 12 or 24μL (Fig. 4B). We fit a logistic regression model to predict rats’ choices on trials where both sides offered guaranteed reward, separately parameterizing each reward volume. The mean regression weight for 6μL was negative, consistent with rats treating this volume as a loss (Fig. 4C).

The win/lose threshold was often reward-history dependent (Fig. 4D). Therefore, we parameterized a dynamic reference point, *r*, as taking one of two values depending on whether the previous trial was rewarded (see Methods). Rewards less than *r* were negative (losses). The relative amplitude of losses versus gains was controlled by the scale parameter *k* (Equation 1; Fig. 1D; fig. S1). Subjective value was reparameterized to include the zero outcome of the gamble, which is a loss when *r* > 0:

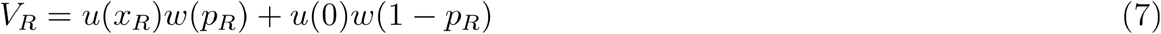

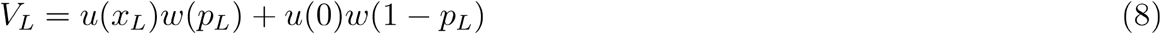

Model comparison using Akaike Information Criteria favored inclusion of the reference point in the model for all rats (fig. S6).

With concave utility, rats should exhibit sharper psychometric performance when the reference point is high (and rewards are more discriminable; Fig. 4E). Indeed, performance was closer to ideal when the reference point was high (Fig. 4F; mean mse between psychometric and ideal performance was 0.143 *low ref* vs. 0.122 *high ref*, p = 3.4e-5, paired t-test across rats).

A loss parameter *k*>1 indicates “loss aversion,” or a greater sensitivity to losses than gains (Equation 1). Kahneman and Tversky (2) estimated a median *k* of 2.25 in humans; we observed a median *k* of 1.66 (Fig. 4G). There was variability across rats, however: 16/36 rats (44%) were not loss averse but were more sensitive to gains (*k*<1). Still, the median *k* across rats suggests, at the aggregate level, a striking similarity to humans (Fig. 4G).

For the first time, we have demonstrated and quantified all of Prospect Theory’s key observations about choice in an animal species; moreover, we did so in rats. Rats exhibit nonlinear (concave) utility for gains, probability distortion, reference dependence, and, frequently, loss aversion. This suggests that the substrates of these phenomena are conserved in rats. Nearly all rats exhibited concave utility, which produced diminishing marginal sensitivity. In Expected Utility Theory, concave utility also indicates risk aversion (9). However in Prospect Theory, concave utility can coincide with risk-seeking behavior due to the elevation of the probability weighting function (21, 22). Indeed, many rats were risk seeking despite showing concave utility functions.

Humans possess two systems for numerical cognition: the approximate number system, which is shared with animals and supports non-symbolic numerical reasoning, and the exact number system, which supports symbolic math and is exclusive to humans (23). Our findings that rats exhibit choice phenomenology from behavioral economics suggests that the approximate number system is sufficient to produce these phenomena, and may contribute to value-based choices across species.

Reward history modified rats’ risk preferences, producing shifts in both the utility and probability weighting functions. While risk preferences are often treated as stable traits, there is evidence that they fluctuate over time and in response to external factors (24). Additionally, we found evidence for a dynamic reference point, in which rats’ treatment of outcomes as gains or losses depended on their reward history. Our data support the notion that choices and preferences are somewhat plastic, flexibly reflecting experience.

Our findings provide a powerful entrypoint for studying the neural circuit basis of choice phenomenology described in behavioral economics, which has wide-ranging impacts on household finances and financial markets.

## Acknowledgements

The authors thank Paul Glimcher, Kenway Louie, Mike Long, David Schneider, Ben Scott, Mikio Aoi, Matthew Lovett-Barron, Cristina Domnisoru, Alejandro Ramirez, and members of the Brody lab for helpful discussions and comments on the manuscript. We thank J. Teran, K. Osorio, L. Teachen, and A. Sirko for animal training. This work was funded in part by a K99/R00 award from NIMH (MH111926, to C.M.C.).

## Author Contributions

All authors provided feedback on analyses and the manuscript. C.M. Constantinople designed and performed all experiments, analyzed the data, and wrote the paper. A.T. Piet provided guidance for modeling and analysis.

## Material and Methods

### Subjects

A total of 39 male rats between the ages of 6 and 24 months were used for this study, including 35 Long-evans and 4 Sprague-Dawley rats (*Rattus norvegicus*). The Long-evans cohort also included LE-Tg (Pvalb-iCre)2Ottc rats (n=5) made at NIDA/NIMH and obtained from the University of Missouri RRRC (transgenic line 0773). These are BAC transgenic rats expressing Cre recombinase in parvalbumin expressing neurons. Animal use procedures were approved by the Princeton University Institutional Animal Care and Use Committee and carried out in accordance with National Institutes of Health standards.

### Behavioral training

Rats were trained in a high-throughput facility using a computerized training protocol. Rats were trained in operant training boxes with three nose ports. When an LED from the center port was illuminated, the animal could initiate a trial by poking his nose in that port; upon trial initiation the center LED turned off. While in the center port, rats were continuously presented with a train of randomly timed clicks from a left speaker and, simultaneously, a different train of clicks from a right speaker. The click trains were generated by Poisson processes with different underlying rates (1, 2); the rates conveyed the water volume baited at each side port. After a variable pre-flash interval ranging from 0 to 350ms, rats were also presented with light flashes from the left and right side ports; the number of flashes conveyed reward probability at each port. Each flash was 20ms in duration; flashes were presented in fixed bins, spaced every 250ms, to avoid perceptual fusion of consecutive flashes (3). After a variable post-flash delay period from 0 to 500ms, the end of the trial was cued by a go sound and the center LED turning back on. The animal was then free to choose the left or right center port, and potentially collect reward.

In this task, the rats were required to reveal their preference between safe and risky rewards. To determine when rats were sufficiently trained to understand the meaning of the cues in the task, we evaluated the “efficiency” of their choices as follows. For each training session, we computed the average expected value per trial of an agent that chose randomly, and a perfect expected value maximizer, or an agent that always chose the side with the greater expected value. We compared the expected value per trial from the rat’s choices relative to these lower and upper bounds. Specifically, the efficiency was calculated as follows:

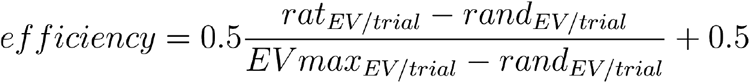

The threshold for analysis was the median performance of all sessions minus 1.5 times the interquartile range of performance across the second half of all sessions. Once performance surpassed this threshold, it was typically stable across months (fig. S7). Occasional days with poor performance were usually due to hardware malfunctions in the rig. Days in which performance was below threshold were excluded from analysis.

### Behavioral model

We fit a behavioral model separately for each rat (see Fig. 1 legend for description of the model). We used Matlab’s constrained minimization function fmincon to minimize the sum of the negative log likelihoods with respect to the model parameters. 20 random seeds were used in the maximum likelihood search for each rat; parameter values with the maximum likelihood of these seeds were deemed the best fit parameters. When evaluating model performance (e.g., Fig. 1E), we performed 5-fold cross-validation and evaluated the predictive power of the model on the held-out test sets.

We initially evaluated three different parametric forms of the probability weighting function, the one- and two-parameter Prelec models and the linear in log-odds model (see below). We compared the different parametric forms using Akaike Information Criteria (AIC), *AIC* = 2*k* + 2*nLL*, where *k* is the number of parameters, and *nLL* is the negative log likelihood of the model. AIC favored the two-parameter Prelec model for nearly all rats, although some rats were equally well-fit by the linear in log-odds model (data not shown). Therefore, we implemented the two-parameter Prelec model.

One-parameter Prelec: 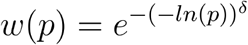

Two-parameter Prelec: 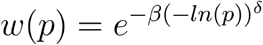

Linear in log-odds: 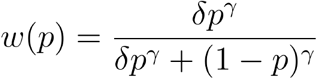

### Psychometric curves

We measured rats’ psychometric performance when choosing between the safe and risky options. For these analyses, we excluded trials where both the left and right side ports offered certain rewards. We binned the data into 11 bins of the difference in the subjective value (inferred from the behavioral model) of the safe minus the risky option. Psychometric plots show the probability that the subjects chose the safe option as a function of this difference (see fig. S2D). We fit a 4-parameter sigmoid of the form:

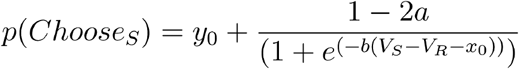

Parameters were fit using a gradient-descent algorithm to minimize the mean square error between the data and the sigmoid, using the sqp algorithm in Matlab’s constrained optimization function fmincon.

### Logistic regression to compare regressors to probability weighting functions

We fit a logistic regression model with a separate regressor for each probability the rat may have been offered (0 to 1 in 0.1 increments), plus a constant term. To compare the regressors to the parametric fits, we normalized the regressors for each probability by subtracting the minimum and dividing by the maximum regressor value, so they ranged from 0 to 1 (Fig. 2G). We computed the mean square error between these normalized regressor values and the probability weighting functions (Fig. 2H). The model was fit using Matlab’s function glmfit.

### Certainty equivalents

#### Non-parametric estimate

We estimated rat’s certainty equivalents by evaluating their psychometric performance (%Chose risky) for each gamble of 48 μL, and estimating the value of the psychometric curve at which performance was at 50% (Fig. 3A). To do this, we fit a line to the two points of the psychometric curve above and below chance level using Matlab’s regress.m function, and interpolated the value of that line that would correspond to 50%.

#### Analytic expression for CE from the model fits

We compared our estimates of rats’ certainty equivalents from their behavioral data to an analytic expression from the subjective probability and utility functions we obtained from the model. We define the certainty equivalent, 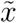, as the guaranteed reward equal to a gamble, *x* with probability *P*. In the case of linear probability weighting, we express this as follows:

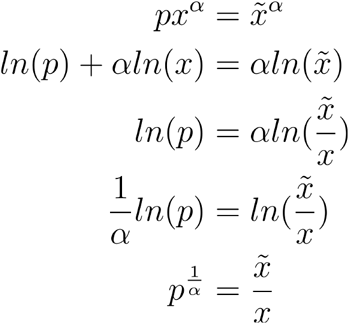

For nonlinear probability weighting, substituting *w*(*p*) for *p* yields an analytic expression for the certainty equivalent from the exponent of the utility function (*α*) and the probability weighting function (also see (4)).

### Behavioral model with reference point

The behavioral model with the reference point (see Fig. 4) was similar to the behavioral model described above, except for elaborations of the subjective utility function *u*(*x*) and subjective value (*V_R_*, *V_L_*). We modified the subjective utility function to include a dynamic reference point, *r*, below which value was treated negatively (as a loss). The relative amplitude of losses versus gains was controlled by the scale parameter *k*.

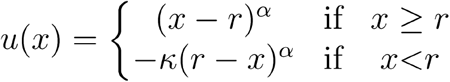

We also reparameterized subjective value. The risky prospect offers two possible outcomes: *x* with probability *P*, and 0 with probability 1 – *p*. In the absence of a reference point, the zero reward outcome (0, 1 – *p*) does not influence choice (0^*α*^ = 0). However, if *r*>0, the zero reward outcome can be perceived as a loss. Therefore, in the reference point model, subjective value was reparameterized to incorporate this possible outcome of the gamble:

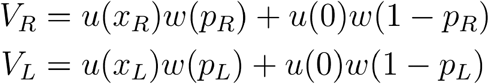

We parameterized the reference point, *r*, to take on two discrete values depending on whether the previous trial was rewarded or not. There were two additional free parameters, *y* and *m* that could account for asymmetric effects of rewarded and unrewarded trials:

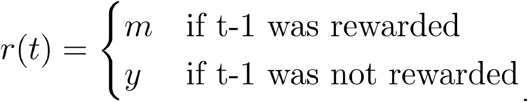

We constrained *r* ≥ 0.

**fig. S1.**
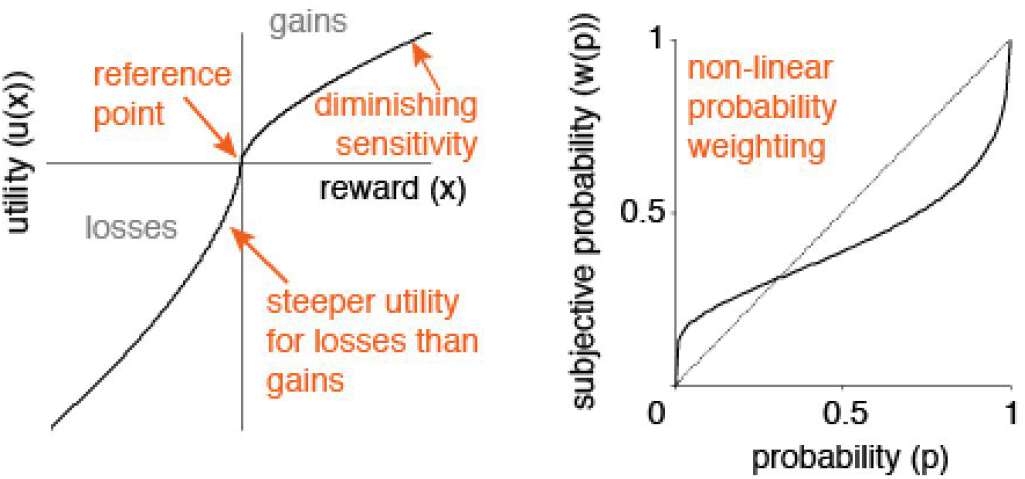
Prospect Theory framework. Prospect Theory describes four aspects of choice behavior that can be understood in terms of the shape of subjective utility and probability weighting functions. Those phenomena, which are schematized here, are (1) diminishing marginal sensitivity (if the exponent of the utility function < 1); (2) loss aversion (the utility function is steeper for losses than gains); (3) reference dependence (rewards are evaluated relative to a reference point); (4) and probability distortion, or nonlinear probability weighting.

**fig. S2.**
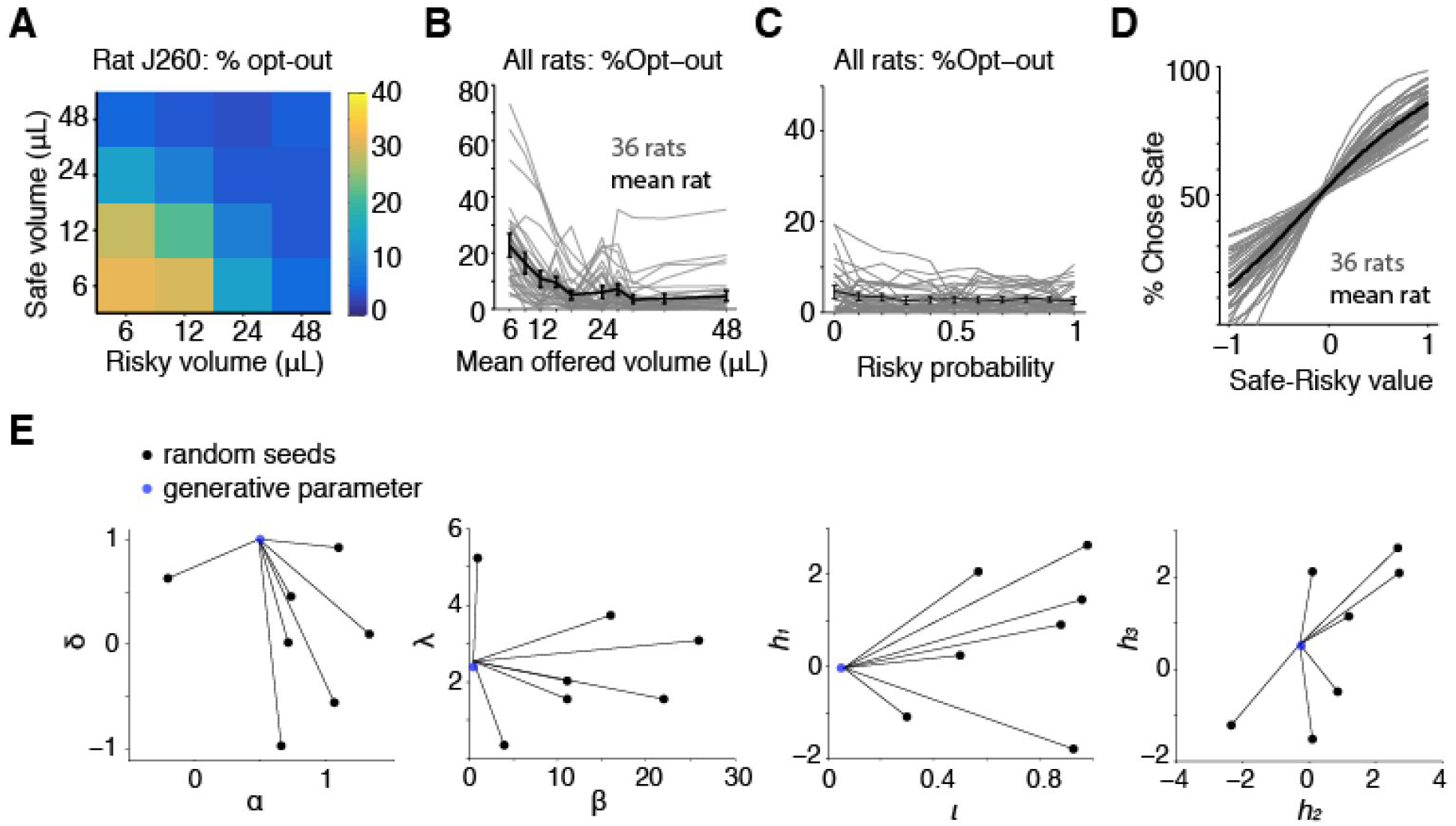
Supplementary analysis of behavioral data and model. **(A)** One rat’s “opt-out” behavior; when both sides offered 6 or 12μL the rat frequently aborted the trial. **(B)** Opt-out behavior for all rats; minimum opt-out rates were subtracted. Black is mean +/− s.e.m. **(C)** Opt-out behavior for all rats as a function of risky reward probability; minimum opt-out rates were subtracted. Black curve is the average +/− s.e.m. across rats. Opt-out behavior did not depend on reward probability. Frequently rats left the center port before all of the flashes were presented, in which case they may not have known which side was risky or safe at the time of opting out. **(D)** Psychometric performance for 36 rats. Gray curves are sigmoid fits (see Methods), black is mean across rats. X-axis is subjective value from model fits. **(E)** Model recovers generative parameters from simulated data. Each plot shows the initial conditions, or random seeds, in black, and the generative parameters, in blue for a pair of parameters in the model. Lines indicate best-fit parameters.

**fig. S3.**
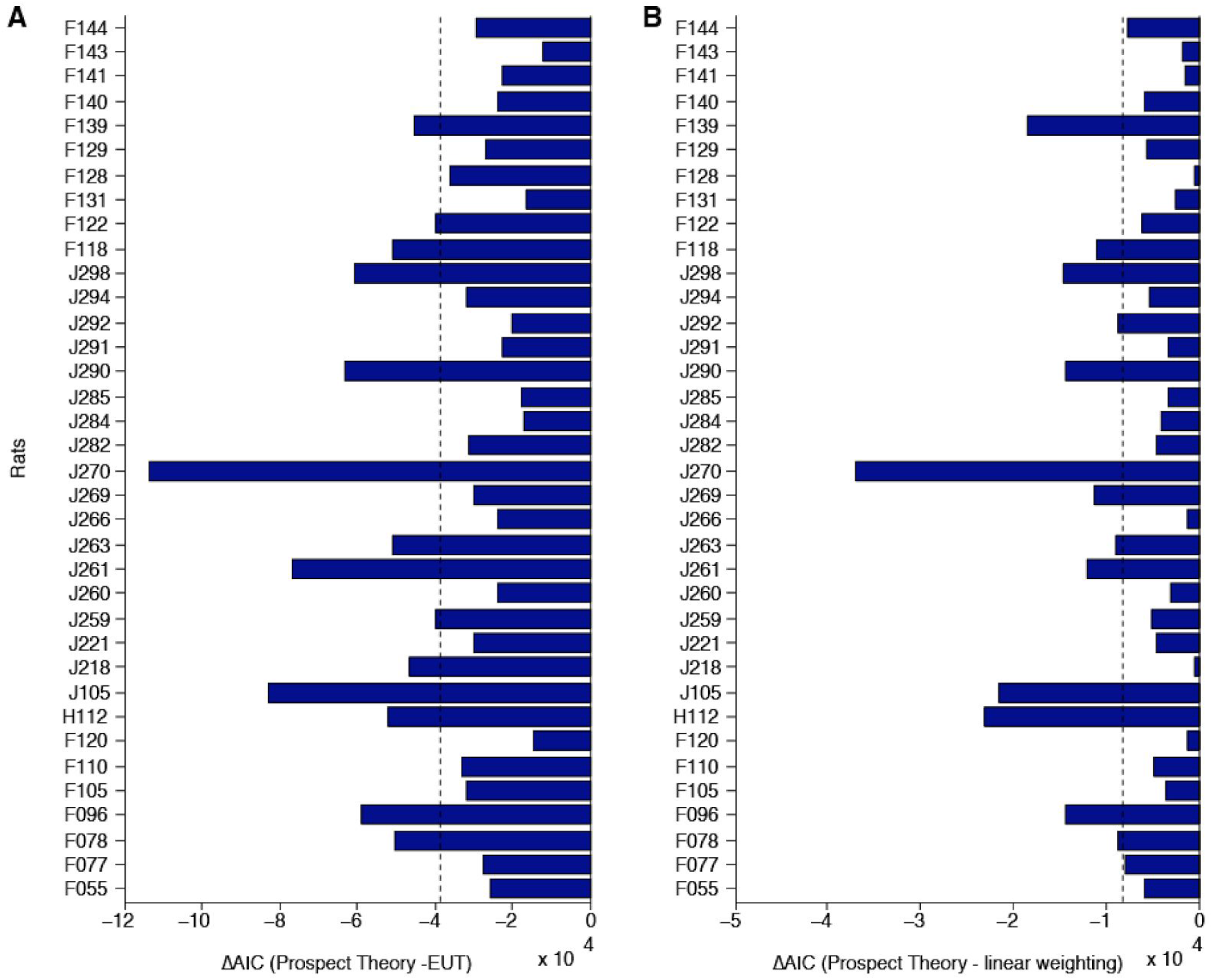
Prospect Theory model outperforms alternative models. **(A)** Difference in AIC values between the Prospect Theory model and the Von Neumann-Morgenstern Expected Utility Theory model (EUT), which is identical to the Prospect Theory model except that it imposes linear probability weighting (5). Negative values indicate the Prospect Theory model was a better fit for each rat. Black dashed line is mean AIC difference. **(B)** Difference in AIC values between the Prospect Theory model and a model that imposed linear weighting of probability and reward volume (the weights applied to each were parameters fit by the model).

**fig. S4.**
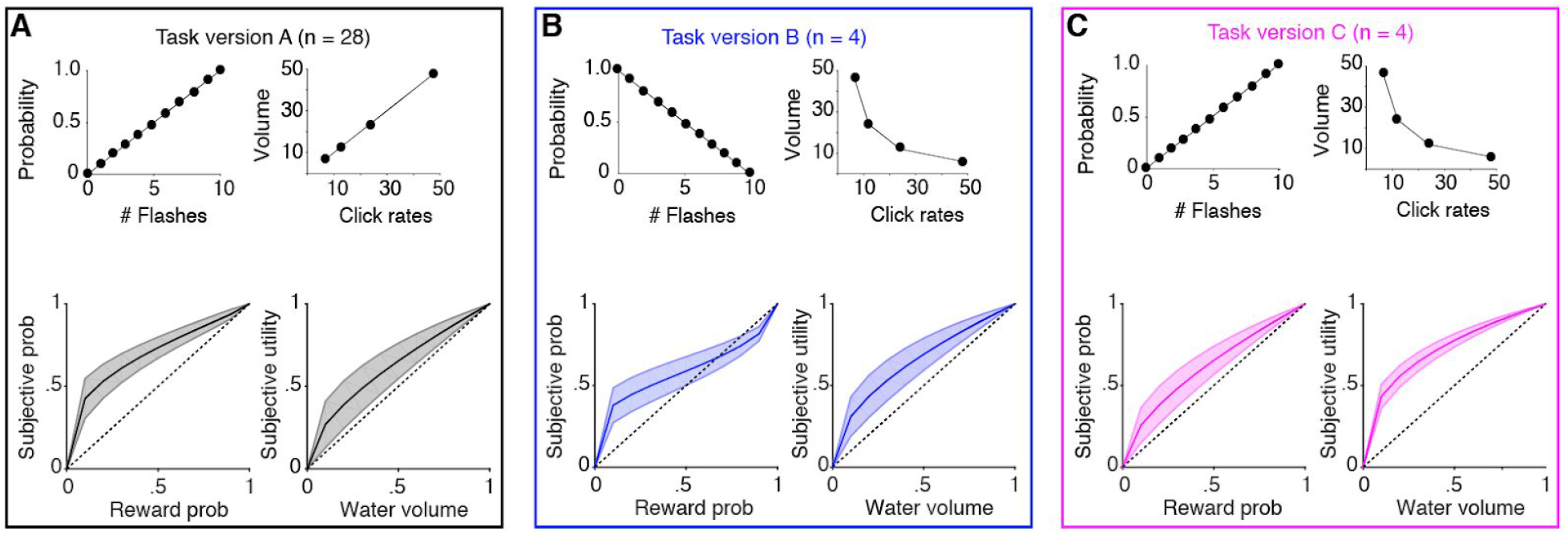
Perceptual noise does not systematically bias model fits. From previous work, we expect the trials with the greatest number of cues (flashes and clicks) to produce the most perceptual noise and, consequently, behavioral variability (1, 2). To determine if perceptual noise on each trial systematically biased the model fits, we trained three cohorts of rats on task versions with different relationships between flashes and reward probability, and clicks and water volume. We reasoned that, if perceptual noise were to systematically influence the model fits, the model fits should look different for rats trained on the different task versions, as they would experience higher perceptual noise on different (and for task versions A and B, precisely opposite) trials. For each task version, we plot the mean +/− s.d. of the probability weighting and utility functions inferred from the behavioral model (bottom panels). On average, rats across task versions all exhibit concave utility (*α* < l) and overweight (especially low) probabilities. **(A)** In task version A, both the flashes and click rates are directly proportional to reward probability and volume, respectively. **(B)** In task version B, the stimuli are inversely proportional to task variables. In this version of the task, the trials with the most perceptual noise (i.e., the highest number of sensory cues) were precisely the opposite of the noisiest trials in task version A. **(C)** In task version C, the relationship between flashes/probability and clicks/volume was incongruent: flashes were directly proportional to probability, and clicks were indirectly proportional to volume.

**fig. S5.**
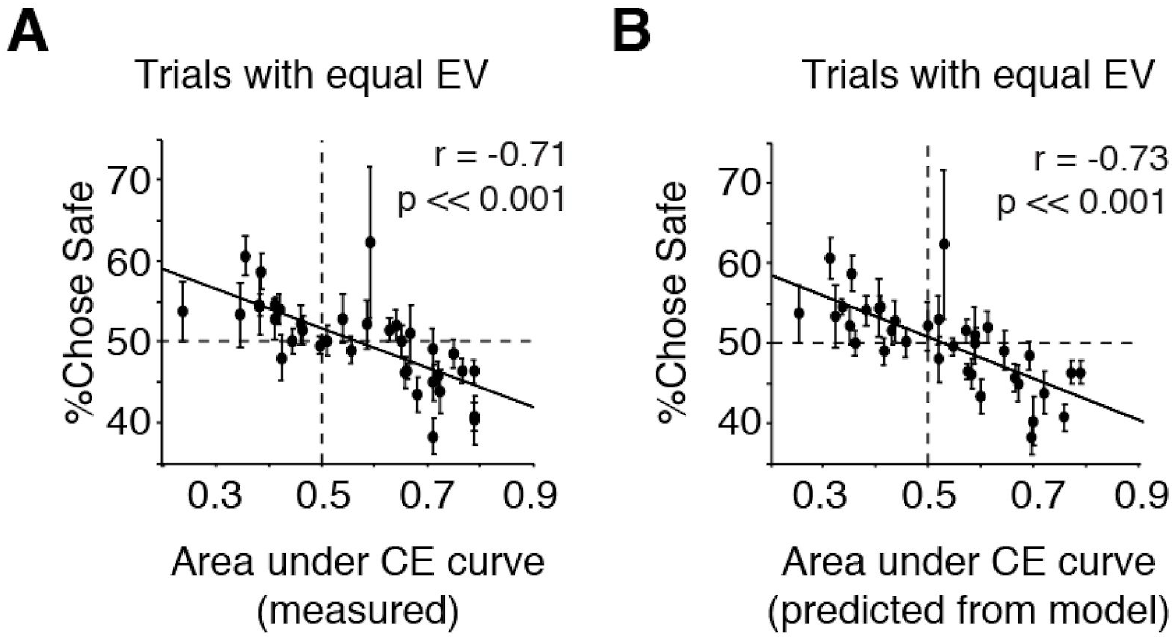
CE area correlates with safe/risky choices on trials with equal expected value. We used the area under the curve of certainty equivalents as a measure of rats’ risk preferences: systematic undervaluing of the gamble (i.e., risk aversion) would produce an area <0.5, and systematic overvaluing the gamble (i.e., risk-seeking) would produce an area >0.5. **(A)** CE area estimated directly from rats’ choices (see Methods) correlates with the probability of choosing the safe option on trials in with safe and risky prospects of equal expected value (mean +/− s.e.m. over days; Pearson’s correlation). Risk-averse rats preferred the safe option on these trials, and risk-seeking rats preferred the gamble. **(B)** Same as A, except CE area was computed using analytic expression from model fits.

**fig. S6.**
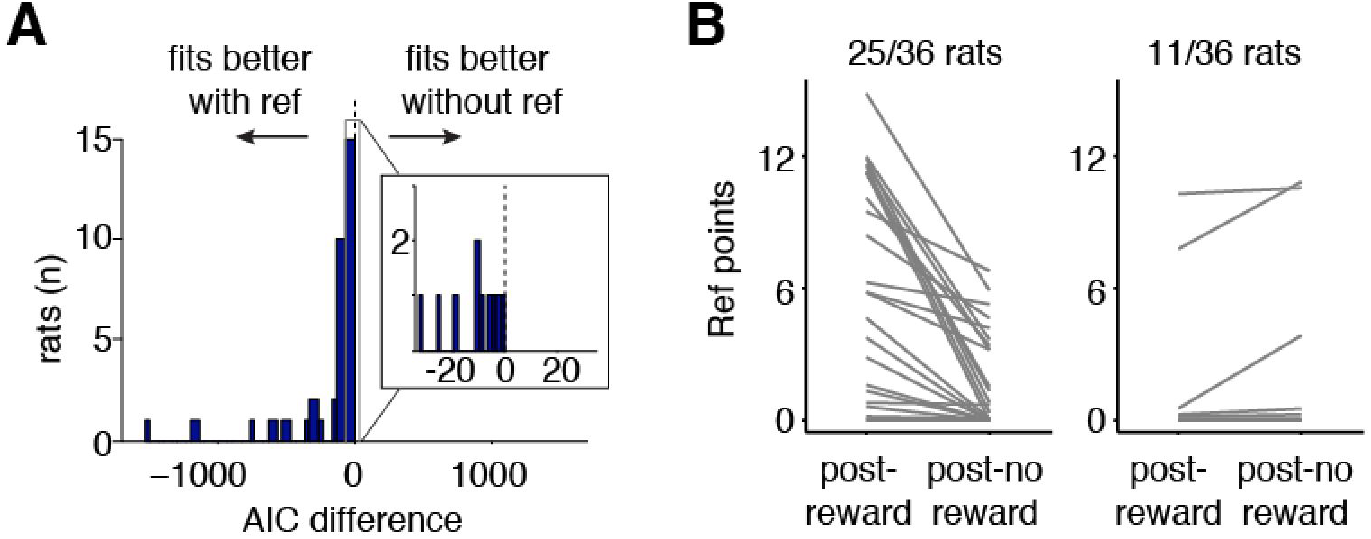
Behavioral model with reference point. **(A)** Model comparison using Akaike Information Criterion (AIC_ref_ - AIC_no ref_) across rats. **(B)** Model estimate of reference points following rewarded and unrewarded trials. Left panel shows rats with higher reference points following rewards. Right panel shows rats with higher reference points following unrewarded trials. Some rats’ reference points were always smaller than 6 but greater than 0, making the zero outcome of the gamble a loss.

**fig. S7.**
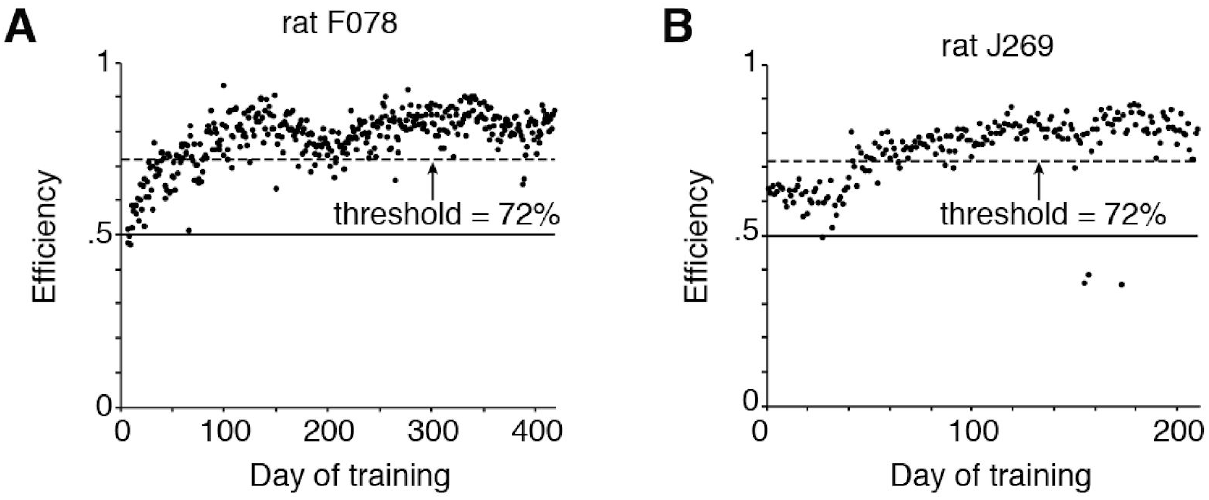
Behavioral performance is stable over days. **(A, B)** Performance (measured in “efficiency”, see Methods) of two rats over days of training. The threshold performance for inclusion in analysis is shown in the dashed lines; for both of these rats it happens to be 72%. Sessions in which performance was below threshold were excluded from analysis.

